# Random Fitness Landscapes are Highly Navigable

**DOI:** 10.64898/2026.05.22.727193

**Authors:** Daniel Oros, Joachim Krug

**Affiliations:** Institute for Biological Physics, University of Cologne, Köln, Germany

## Abstract

With the increasing availability of large scale empirical fitness landscape data, there is a need for simple yet informative null models that can be used to interpret metrics of landscape ruggedness and navigability. A natural choice of a null model that maximizes ruggedness in a statistical sense assigns independent and identically distributed fitness values to the genotypes, a setting often referred to as the House-of-Cards (HoC) or mutational landscape model. In this work we examine the navigability of these landscapes, as quantified by the mean size of the adaptive basins of local fitness peaks. The adaptive basin is the set of genotypes from which a peak can be reached via selectively accessible, i.e., strictly fitness-increasing mutational paths. Building on recent rigorous results on the statistics of accessible paths, we show that the adaptive basins in the HoC landscape encompass a positive fraction of all genotypes that is an analytically computable, increasing function of the number of alleles per site. For the four letter nucleotide alphabet, an average peak basin contains 52.8 % of all genotypes. When conditioned on peak fitness, the expected basin size increases linearly with fitness rank. The exact results on adaptive basins are complemented by an approximate analysis of gradient basins formed by greedy adaptive paths which maximize the fitness increase in each step. We argue that recent reports of large adaptive basins in empirical fitness landscapes should be reinterpreted in the light of our findings.

## Introduction

Fitness landscapes have been studied extensively since their introduction almost a century ago by Wright (1932). In its simplest form, the fitness landscape maps the genotype of an organism to its reproductive success, denoted by fitness, in a given environment. While being a dramatic simplification of the complex biology underlying the evolutionary process, the concept is useful for addressing broader conceptual questions, such as the patterns of repeatability and predictability of evolutionary trajectories (Svensson and Calsbeek, 2012; Lobkovsky and Koonin, 2012; De Visser and Krug, 2014; Srivastava et al., 2026) and the way these are shaped by epistasis (Phillips, 2008; Starr and Thornton, 2016; Domingo et al., 2019; Bank, 2022). When the time between mutations is much larger than the fixation time, one can think of a genetically homogeneous population moving across the landscape one mutational step at a time (Gillespie, 1984). The fitness value increases at each step and the population eventually reaches a fitness peak, a genotype having larger fitness than all its single mutant neighbors. Following Stadler and Stadler (2010), we distinguish between adaptive paths, where every beneficial mutation can fix no matter how small the fitness gain, and gradient or greedy paths, where in each step the largest fitness increase achievable by a single mutation fixes.

The past two decades have seen a steady and rapid increase in the scale of the empirical fitness landscapes that have been explored experimentally (Weinreich et al., 2006; Szendro et al., 2013; De Visser and Krug, 2014; Bank et al., 2016; Aguilar-Rodríguez et al., 2017; Pokusaeva et al., 2019), and in the most recent studies tens to hundreds of thousand viable genotypes were characterized in terms of fitness or some proxy thereof (Moulana et al., 2022; Papkou et al., 2023; Westmann et al., 2024; Antunes Westmann et al., 2024; Chattopadhyay et al., 2025; Gaszek et al., 2025). Some of these works find fitness landscapes that are highly rugged yet their peaks are highly accessible, and it has been argued that this high navigability can not be explained by current fitness landscape models (Papkou et al., 2023). Here we examine the relationship between landscape ruggedness and peak accessibility within the framework of the *House of Cards* (HoC) model (Kingman, 1978; Kauffman and Levin, 1987; Jain and Krug, 2007), also referred to as the mutational landscape model (Gillespie, 1984; Orr, 2002; Rokyta et al., 2005), which assigns fitness values to genotypes as independent random variables drawn from a continuous fitness distribution. The HoC fitness landscape is maximally rugged in statistical sense (Szendro et al., 2013), and as such it constitutes a natural choice as a null model for comparison to empirical landscapes. Our main finding is that, even within this extremely simplified setup, high ruggedness and high navigability are not mutually exclusive.

A large number of previous studies have investigated various aspects of the HoC model. These include the length of adaptive walks until a peak is reached, for adaptive (Kauffman and Levin, 1987; Macken and Perelson, 1989; Orr, 2002; Neidhart and Krug, 2011) and gradient walks (Orr, 2003), as well as the statistics of fitness-increasing accessible paths (Weinreich et al., 2005; Franke et al., 2011; Hegarty and Martinsson, 2014; Zagorski et al., 2016; Berestycki et al., 2017; Schmiegelt and Krug, 2023). Here we focus on the size of the basins of attraction of fitness peaks, defined as the number of non-peak genotypes from which the peak can be accessed via adaptive or gradient paths (Stadler and Stadler, 2010). To our knowledge, analytical results for adaptive basin sizes in probabilistic fitness landscape models have not been reported previously, apart from lower bounds for certain models of structured landscapes with global constraints (Das et al., 2020; Krug and Oros, 2024; Pahujani and Krug, 2025). We provide an exact expression for the expected adaptive basin size of the HoC model in the limit of long genotype sequences and show numerically that our solutions perform well already for moderate sequence lengths. Genotypes of high fitness and especially peaks are accessible from a large fraction of the whole fitness landscape, making HoC landscapes highly navigable under adaptive dynamics. By contrast, an approximate analysis shows that the basins formed by gradient paths are small, making HoC landscapes much less navigable under greedy dynamics.

We argue that the notion that rugged landscapes are generally poorly navigable stems from either considering only direct paths as possible adaptive trajectories (Franke et al., 2011; Hegarty and Martinsson, 2014), or from the commonly used low-dimensional metaphor of a mountainous terrain. We allow for indirect paths and study the HoC landscape mathematically rather than relying on low-dimensional intuition. In this way we elucidate how the accessibility of peaks and the navigability of the landscape emerge as the genotype space grows. Our results can be viewed as a conservative lower bound for accessibility of real fitness landscapes as they are derived in a maximally rugged setting.

## Model

We define a genotype as a string of *L* loci, where each locus takes on one of *A* different values. The genotype space is composed of *A*^*L*^ genotypes, and the number of one-mutant neighbors of a genotype is *n* = *L*(*A*− 1). Referring to distances between two genotypes will be done interchangeably by their Hamming distance *d* or their re-scaled Hamming distance 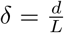, depending on context. Mathematically speaking, the genotype space forms a Hamming graph (Stadler, 2002) on which a monomorphic population characterized by its genotype moves one mutational step at a time. The restriction to fitness-increasing adaptive steps orients the links of the Hamming graph in the direction of higher fitness, which gives rise to an oriented acyclic structure known as a fitness graph (De Visser et al., 2009; Crona et al., 2013, 2017).

We follow the definition of Stadler and Stadler (2010) for gradient and adaptive basins. A basin is associated with a target genotype that will usually be a fitness peak, though it will turn out to be useful to apply the concept to non-peaks as well. By convention, the target genotype is part of its basin, which implies that the minimal basin size is 1. The *adaptive basin* of a genotype *σ* is composed of all genotypes from which *σ* can be accessed via a fitness-increasing path. Paths can be direct, where the Hamming distance to the target decreases in each step, or indirect (Krug and Oros, 2024). In the latter case the number of mutational steps along the path exceeds the Hamming distance to the target. The *gradient basin* of a genotype is composed of all genotypes that can reach the target by following the mutations that confer the highest fitness gain in their single-mutant neighborhood. The union of all gradient basins of all fitness peaks comprises all genotypes in the full landscape, as each genotype belongs to the basin of a single fitness peak. This however is not true for adaptive basins. Figure 1**a** exemplifies both basin types in a small landscape.

**Figure 1.**
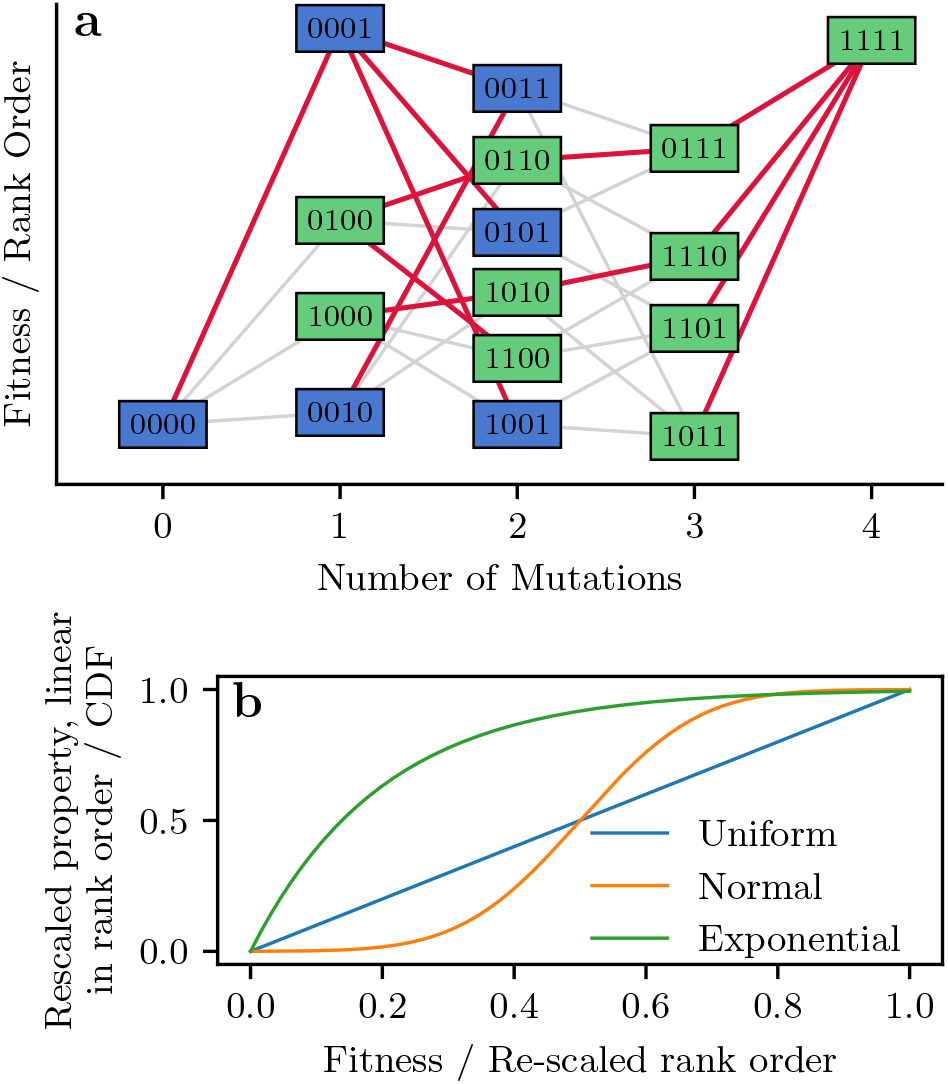
Fitness landscape and fitness rank. a Schematic of a fitness landscape for *L* = 4 and *A* = 2. Lines connect nearest neighbors and the highest fitness increase for each genotype is marked in red. Genotypes 0001 and 1111 are fitness peaks. Their respective adaptive basins consist of 15 and 14 genotypes, because 0011 is not contained in the basin of 1111. The corresponding gradient basins are marked in blue and green. b Cumulative distribution function for uniform *U* (0, 1), exponential Exp(5) and normal 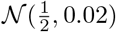 fitness distributions. Uniform fitness is proportional to rank order, whereas the other distributions introduce a nonlinear relation between rank and fitness.

Here we study the structure of adaptive and gradient basins in the HoC model, where genotypes are assigned independent, identically distributed fitness values. This choice reflects a null model with no correlation between fitness values, which generates landscapes that are maximally rough in a statistical sense compared to single peaked landscapes (Szendro et al., 2013). We do not allow for regions of neutrality, but argue that these would only increase the adaptive basin size of genotypes if evolution along neutral paths is permitted (Greenbury et al., 2022). In fact, we make the somewhat stronger assumption that no two genotypes have exactly the same fitness, which implies that the landscape possesses a unique global rank order of fitness values. This is realized naturally by choosing the HoC fitness values from a continuous probability distribution.

Since peaks, accessible paths and gradient as well as adaptive basins are fully defined by the rank ordering of the fitness values, they are not altered by strictly monotonic transformations of fitness. As a consequence, provided it is continuous, the choice of the underlying fitness distribution is irrelevant for the properties of interest in this work. For convenience, in the following we choose the fitness values to be uniformly distributed in the unit interval (0, 1] To motivate this choice, let *R*_*σ*_ be the rank of genotype *σ* between 1 (lowest fitness) and *A*^*L*^ (highest fitness) in the global rank order 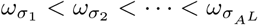 . Then the cumulative distribution function *F*_*U*_ (*ω*_*σ*_) = *ω*_*σ*_ of the uniform distribution evaluated for a genotype *σ* converges to the re-scaled rank 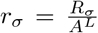 for large *A*^*L*^. Figure 1**b** show examples of different fitness distributions and illustrates how a property that is linear in rank gets distorted when using fitness instead of rank.

To quantify the navigability of the HoC landscape, we adopt and modify a definition introduced by Greenbury et al. (2022) in the context of genotype-phenotype-fitness maps. For a pair of genotypes *σ* and *τ*, let Ψ(*τ, σ*) be 1 if there is a fitness increasing path from *τ* to *σ* and 0 if there isn’t. By convention Ψ(*σ, σ*) = 1. Then the *peak navigability* of the landscape is defined as the average of Ψ over all pairs of genotypes (*τ, σ*_*P*_) where the target is a peak,

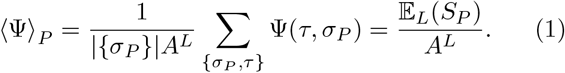

Here ∣{ σ_*P*_}∣ is the number if peaks

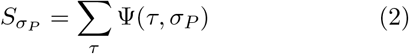

denotes the size of the adaptive basin of peak *σ*_*P*_ with expectation value 𝔼_*L*_(*S*_*P*_), where we use the subscript *L* to indicate the dependence of expectation values on the sequence length. By construction ⟨Ψ⟩ _*P*_ lies between 0 and 1. We say that a landscape is *highly navigable* if ⟨Ψ⟩ _*P*_ converges to a positive value for large *L*, which implies that the adaptive basin of a peak on average contains a positive fraction of all genotypes.

## Results

In the following we present our main results. Detailed derivations can be found in the Appendices.

## Adaptive Basins

Our analytical results for the adaptive basins are based on the mathematical theory of *accessibility percolation*, which is concerned with the statistical properties of the random variables Ψ(*τ, σ*) that encode the existence of fitness-increasing paths between genotypes *τ* and *σ* (Nowak and Krug, 2013; Krug, 2021; Schmiegelt and Krug, 2023). The key result of the theory is that, for genotype pairs (*τ*_*L*_, *σ*_*L*_) at distance *d*(*τ*_*L*_, *σ*_*L*_) ∼ *L*, in the limit *L* → ∞ the probability of existence of a path transitions abruptly from 0 to 1 when the fitness difference 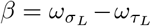 exceeds a threshold value *β*_*c*_ that depends only on the rescaled distance 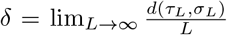 and the number of alleles *A*. The threshold *β*_*c*_(*δ, A*) is given by the unique root in *β* of the function (Schmiegelt and Krug, 2023)

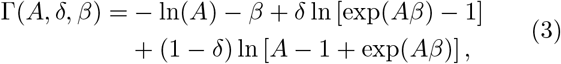

provided this root lies within the unit interval.

To estimate the adaptive basin size of a target genotype of fitness *ω*, we note that from any genotype there are

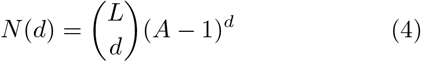

genotypes at distance *d*. For large *L, N* (*d*) becomes sharply peaked at the rescaled distance 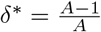 . Thus the vast majority of genotypes contributing to the adaptive basin of the target are located at this distance, and an accessible path from such a genotype to the target exists with high probability if its fitness is below *ω* −*β*^*\**^, where *β*^*\**^(*A*) = *β*_*c*_(*δ*^*\**^, *A*). Under the uniform fitness distribution this is true with probability *ω*− *β*^*\**^ if *ω > β*^*\**^. By linearity of expectation, the expected value of the adaptive basin size *S*(*ω*) of the target is therefore given by

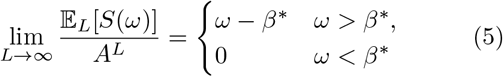

asymptotically for large *L*. Thus for genotypes with fitness above the threshold *β*^*\**^, the adaptive basin is predicted to contain a positive fraction of all genotypes that increases linearly with fitness on the uniform scale, i.e., with relative fitness rank. The fitness of a peak genotype exceeds that of its *n* neighbors, and therefore the probability density of the peak fitness is

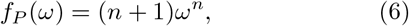

which is concentrated near *ω* ≈ 1 for large *L*. This results in the mean adaptive basin size for peaks to be

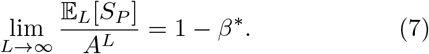

To determine *β*^*\**^ one can transform Eq. (3) into a polynomial of degree *A* (Schmiegelt and Krug, 2023), see table 1 for analytical solutions for *A* = 2, 3 and 4. For the canonical case *A* = 4 corresponding to the nucleotide alphabet, *β*^*\**^≈ 0.472, which implies according to Eq. (7) that the adaptive basin of a peak typically contains about 52, 8% of all genotypes. Performing a large *A* expansion of Eq. (3) for *δ* = *δ*^*\**^ results in

**Table 1.** Exact values for *β*^*\**^(*A*) for *A* = 2, 3 and 4.

| $A$ | $\beta^*$ |
| --- | --- |
| 2 | $\frac{1}{2} \ln(2 + \sqrt{5}) \approx 0.722$ |
| 3 | $\frac{1}{3} \ln\left(2\sqrt{10} \cos\left(\frac{1}{3} \arccos\left(-\frac{1}{10\sqrt{10}}\right)\right)\right) \approx 0.565$ |
| 4 | $\frac{1}{4} \ln\left(\sqrt{1+x} + \sqrt{\left(2-x + \frac{62}{\sqrt{1+x}}\right)}\right) \approx 0.472$<br>with $x = 4 \cdot \sqrt[3]{15}$ |

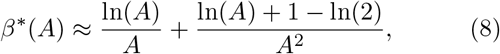

which is a decreasing function of *A*. Thus the average size of peak adaptive basins increases with the number of alleles, reaching 84% of all genotypes for *A* = 20. Figure 2**a** compares the expansion Eq. (8) to numerically determined values of *β*^*\**^ up to *A* = 20.

**Figure 2.**
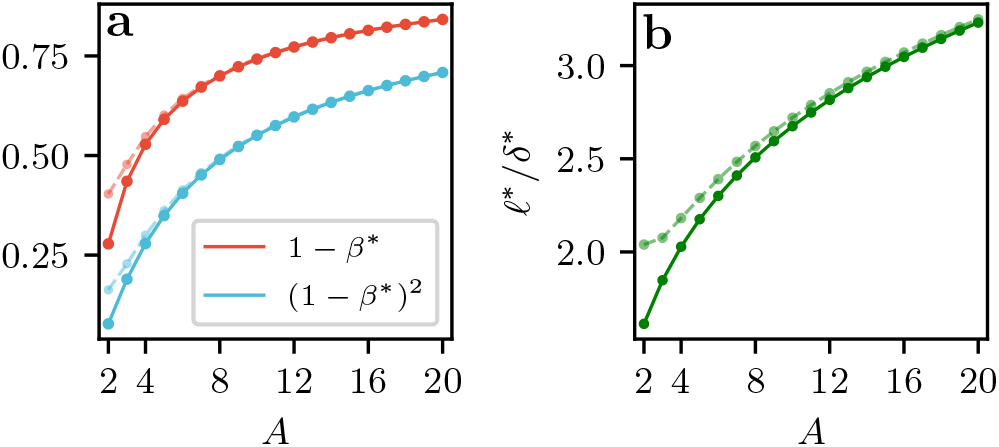
Navigability and adaptive path length increase with the number of alleles. a Upper line shows the predicted fraction of all genotypes contained in an average adaptive basin of a peak [Eq. (7)], lower line shows the predicted pairwise navigability [Eq. (10)]. b Predicted excess length of adaptive paths at the ac-cessibility threshold. In both panels, solid lines show values obtained by numerically finding the exact root *β*^*\**^ of the function Γ(*A, δ, β*) defined in Eq. (3) at *δ* = *δ*^*\**^, and dashed lines make use of the large *A* approximation [Eq. (8)].

Figure 3 displays the results of numerical simulations for the adaptive basin size in comparison to the asymptotic predictions of Eqs. (5) and (7). The finite *L* data for the average of the fitness-dependent basin size *S*(*ω*) in Fig. 3**a** show some rounding of the sharp transition at *ω* = *β*^*\**^, in particular for small *A*, but align nicely to the predicted line for larger *ω*. The data for the mean basin size of peaks *S*_*P*_ lie above the asymptotic prediction for small *L* and approach it slowly from below for large *L* (Fig. 3**b** and **c**). The slow convergence is expected, since the leading order corrections to the threshold fitness difference *β*_*c*_ are of order 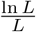 and the change in probability of existence of paths occurs over a range of order (Schmiegelt and Krug, 2023). Additional 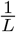-corrections arise in the derivation of the expected basin size, see Appendix B. Extending the numerical simulations to larger of *L* and *A* is challenging, as the numerical complexity of calculating adaptive basins in the fitness graphs with Hamming graph structure is 𝒪 (*A*^*L*+1^*L)* (Cormen et al., 2022). Nevertheless the simulations provide strong evidence for the validity of our asymptotic predictions.

**Figure 3.**
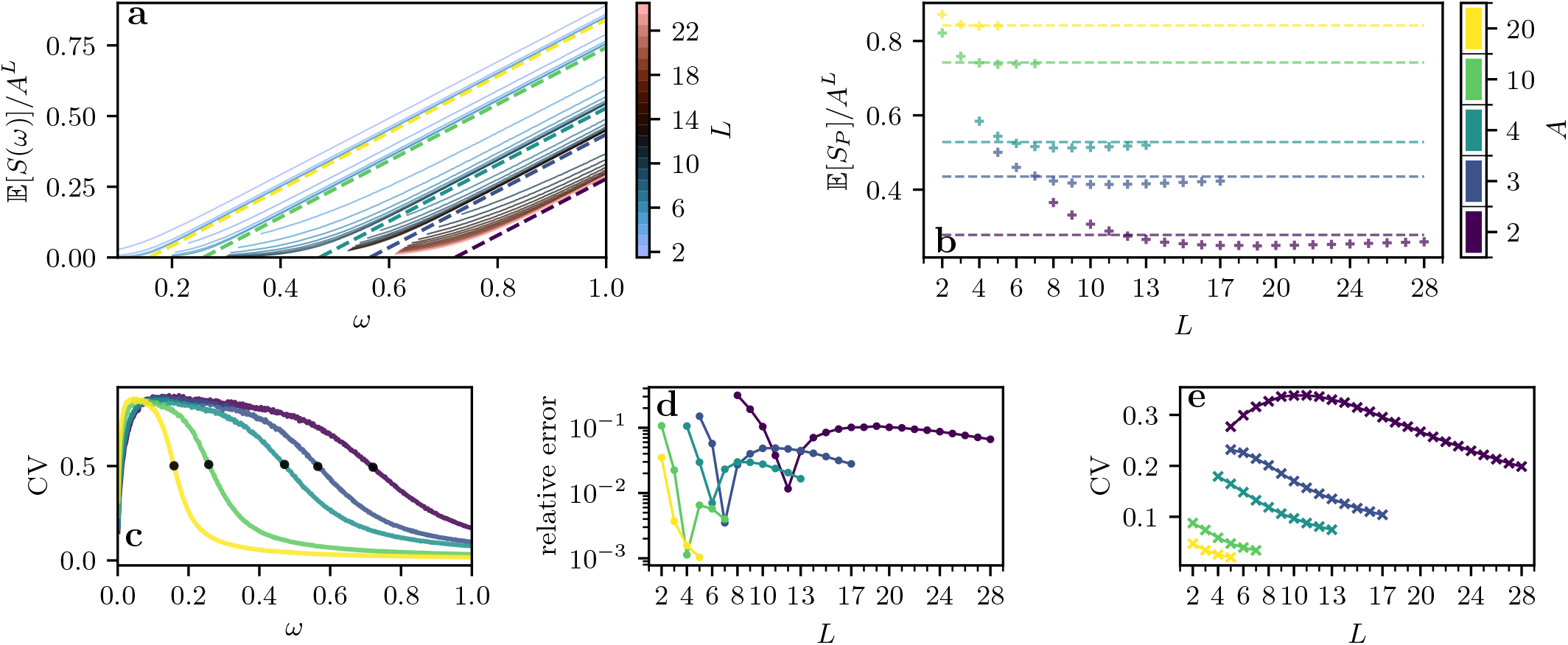
Adaptive basin size. a Dashed straight lines are the analytical prediction Eq. (5) for the asymptotic mean adaptive basin size as a function of the target fitness *ω* for different value of *A*, encoded by the color bar in panel b. The results of numerical simulations for non-peak genotypes are shown as solid lines for different values of *L*, encoded by the color bar in panel a, and cut-off early for better visibility. b Mean adaptive basin size of peaks as a function of *L* for different values of *A*. The figure compares the results of numerical simulations to the asymptotic prediction Eq. (7) shown as dashed horizontal lines. c Coefficient of variation (CV) for the largest simulated value of *L* for the corresponding data shown in panel a. The CV generally decreases with increasing *A* after the predicted fitness threshold *β*^*\**^ marked on the individual curves with black dots. d Relative error of the numerical values from b with respect to the asymptotic prediction Eq. (7). The dip in the data marks the point where the numerical data cross the asymptotic prediction. The error is a decreasing function of *A* and displays a slow decline for large *L*. e Coefficient of variation for the numerical mean shown in panel b and additional data points for *A* = 2 for small *L*. The CV decreases with increasing *A* and increasing *L* for large *L*.

### Pairwise navigability and path length

Figure 2 displays two further quantities of interest that can be extracted from the analysis of Schmiegelt and Krug (2023). First, consider the fraction of all pairs of genotypes for which there exists an accessible path in either direction from one to the other, which defines the *pairwise navigability*

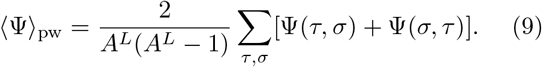

For large *L*, the re-scaled distance between the genotypes is close to *δ*^*\**^ with high probability, and ⟨Ψ⟩_pw_ can therefore be approximated by twice the area under the curve from Eq. (5), which yields

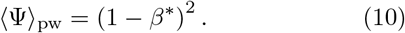

The lower curve in figure2**a** compares numerically exact values of this quantity to the large *A* expansion of Eq. (8).

Furthermore, Theorem 3 of Schmiegelt and Krug (2023) states that the expected re-scaled length *ℓ*^*\**^ of an accessible path between two genotypes with fitness difference *β*^*\**^ at distance *δ*^*\**^ is given by

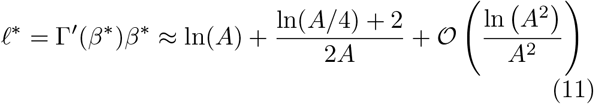

for *L*→ ∞, where the approximation uses Eq. (8) for *β*^*\**^. This is the path length between genotypes at distance *δ*^*\**^ with minimal fitness difference *β*^*\**^ for which an accessible path exists. Increasing the fitness difference increases the number of potential paths, which is expected to lower the average path length. As a con-sequence, *𝓁*^*\**^ can be interpreted as an *upper bound* for the re-scaled length of paths contributing to the adaptive basin. Figure 2**b** shows the *excess length 𝓁*^*\**^*/δ*^*\**^ as a function of *A*, which quantifies the fraction of indirect, non-distance decreasing steps along the path. The excess length, and thus the contribution of indirect paths to the adaptive basins, increases monotonically with *A*.

### Gradient Basins

The scale of the size of gradient basins is set by the number of neighbors *n* in the genotype graph. The mean gradient basin size of peaks can be obtained from a simple counting argument (Franke and Krug, 2012). Noting that each genotype belongs to the gradient basin of exactly one peak, and the expectation value of the number of peaks *N*_*P*_ is (Kauffman and Levin, 1987)

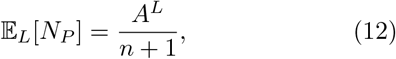

the expecation value of the gradient basin size of a peak *G*_*P*_ can be estimated as

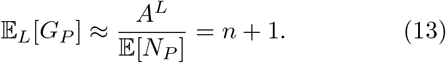

Since *N*_*P*_ is asymptotically normal with a variance that is of the order of the mean (Macken and Perelson, 1989; Baldi and Rinott, 1989), this estimate is exact up to small corrections of order 𝒪 (*A*^*−L/*2^). It is interesting to note that *n* + 1 is the *minimal* size of the adaptive basin of a peak. In contrast, gradient basins may contain only the focal genotype. On the other hand, as pointed out by Franke and Krug (2012), the gradient basin of the global fitness maximum is at least of size *n* + 1.

In the Appendix we show that the probability of a direct, strictly fitness increasing gradient path of length *d* from a genotype with fitness *ω*_0_ to one with fitness *ω* can be approximated by

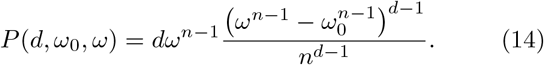

We neglect non-direct paths as they are exponentially less likely than direct ones. Starting genotypes have by definition fitness *ω*_0_ *<* 1, allowing to set 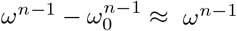 for large *n*. Summing over all genotypes grouped by their Hamming distance *d* from the focal genotype then results in the expression

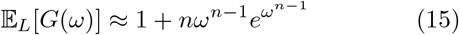

for the expected size of the gradient basin of a (peak or non-peak) genotype of fitness *ω*. Integrating (15) over the peak fitness distribution (6) recovers (13) to leading order in *n*. As the gradient basin size grows only linearly in *L*, simulations can be extended to larger systems than for adaptive basins, see Fig. 4 for a comparison of the theoretical prediction and numerical results.

**Figure 4.**
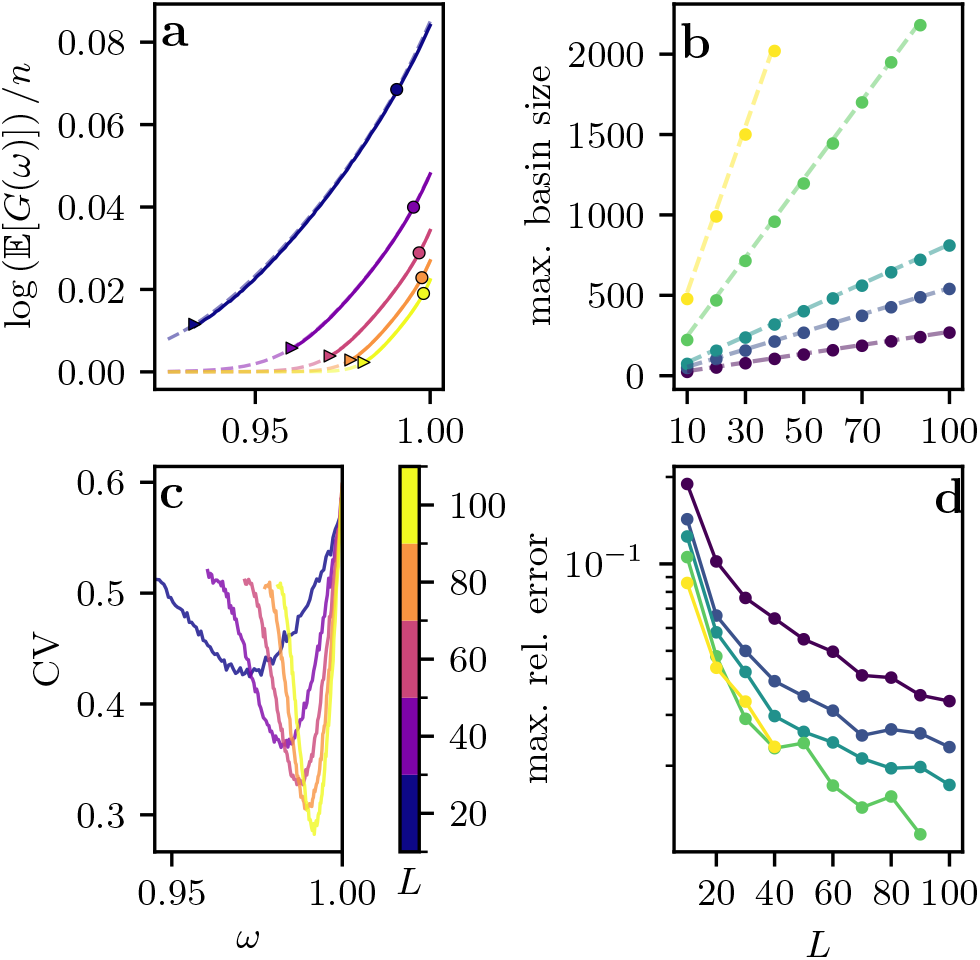
Gradient basin size. a Mean gradient basin size for *A* = 4 and selected values of *L* as a function of the target fitness *ω*. Dashed lines are the analytical approximation from Eq. (15) and solid lines are numerical simulations. The scaling is chosen to highlight how well the numerical simulations agree with the approximate analytical calculation. Dots show the mean gradient basin size of peaks *n*+1 [Eq. (13)] and the triangular markers highlight where Eq. (15) is equal to 2. b Numerical mean gradient basin size for genotypes of fitness close to *ω* = 1 are shown as dots, and the analytical approximation *ne* + 1 obtained from Eq. (15) is shown as dashed lines. c CV values for the data shown in a, illustrating that the distribution is most sharply concentrated for basins of intermediate mean size. d Maximal relative error of the mean gradient basin size for different values of *A* as a conservative measure for how well the analytical solution approximates the simulations. Convergence for large *L* is apparent from the figure. Panels b and d use the same color bar for *A* as in Fig.3.

Equation (15) predicts that the largest gradient basins of genotypes with maximal fitness *ω* = 1 are of size *ne* + 1, exceeding the mean size Eq. (13) by a factor of *e* (Fig. 4**b**). With decreasing target fitness the expected gradient basin size decreases rapidly. Only the highest fitness genotypes have gradient basins that contain more genotypes than the target itself. In Fig. 4**a** we illustrate this for *A* = 4 by depicting the fitness values *ω*_2_ at which the expected basin size 𝔼 [*G*(*ω*_2_)] = 2, showing that *ω*_2_ tends to unity with increasing *L*. Figure 4**c** shows the corresponding CV values, which are smallest for moderate sized gradient basin sizes between fitness *ω*_2_ and 1. The same pattern is found for other values of *A*. Figure 4**d** shows the maximal relative error, for fitness values between *ω*_2_ and 1, as a conservative measure for the validity of our theoretical prediction Eq. (15) compared to numerical simulations. These data show that the tree-like approximation we used to derive Eq. (15) becomes more accurate for larger *L*.

## Discussion

### Adaptive basins are larger than expected

Our analysis shows that typical adaptive basins in uncorrelated random HoC fitness landscapes contain a positive fraction of all genotypes, which increases with the number of alleles *A* and reaches 84 % for *A* = 20, corresponding to the amino acid alphabet. This finding challenges the established view that landscape ruggedness restricts navigability. The structure and navigability of a fitness landscape is determined primarily by the strength and distribution of pairwise sign-epistatic interactions, where the sign of a mutational effect at a locus depends on the state of other loci (Weinreich et al., 2005; Poelwijk et al., 2007; Srivastava et al., 2026). A closer look at the dual effects of sign-epistatic interactions may therefore help to resolve this apparent paradox.

Pairwise sign-epistasis can be simple or reciprocal, where either one or both mutations reverse each other’s effects respectively. The HoC model displays high levels of sign epistasis of both types, with 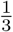 of two-way interactions showing simple and reciprocal sign epistasis each (Szendro et al., 2013). Reciprocal sign epistasis is a necessary condition for multi-peaked landscapes (Poelwijk et al., 2011; Riehl et al., 2022; Saona et al., 2022), and both types of sign epistasis reduce the number of direct accessible paths to a fitness peak (Weinreich et al., 2005; Bank, 2022). However, sign epistasis also creates evolutionary detours, indirect paths with mutational reversals or sideways steps, where a resident allele is replaced by another allele that is not present in the target sequence (Wu et al., 2016; Zagorski et al., 2016; Ribeca et al., 2026).

The decreasing probability of finding a fitness monotonic sequence of fitness values in the rugged landscape is therefore offset by the increase in the number of potential indirect paths. As expected, this effect becomes more pronounced with increasing alphabet size. Our theoretical results predict, supported by simulations, that with increasing *A* the fraction of genotypes that do *not* belong to the adaptive basin of a typical peak decreases 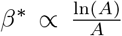 [Eqs. (7, 8)], whereas the excess length of a critical accessible path increases as *𝓁*^*\**^*/δ*^*\**^ ∼ ln *A* [Eq. (11)]. In this view, the intuition responsible for underestimating the adaptive basin sizes in the HoC landscapes derives from an inappropriate focus on direct accessible paths.

Another confounding factor affecting the intuition about the navigability of the HoC landscape is the failure to clearly distinguish between adaptive paths and adaptive walks. The accessible adaptive paths considered in this study are features of the fitness landscape itself, which can be defined and studied without reference to any particular evolutionary dynamics. In contrast, adaptive walks implement a specific dynamic rule where a monomorphic population moves to a neighboring genotype with higher fitness (Kauffman and Levin, 1987; Macken and Perelson, 1989; Orr, 2002). Although every monotonic fitness increasing path is accessible and therefore a valid trajectory of an adaptive walk, this does not mean that the walker has a high chance of finding those paths. Macken and Perelson (1989) cal-culated the mean path length for a random adaptive walker to reach a fitness peak in the HoC landscape to be ∝ ln(*n*) ∼ ln(*L*), with a variance of the same order. The weight that the adaptive walk dynamics assigns to paths of length 𝒪 (*L*) that dominate the adaptive basins is therefore very small. The problem of quantifying the *reachability* of fitness peaks, defined as the fraction of adaptive walkers that reach the peak from a uniformly chosen starting point (Das et al., 2020; Li and Zhang, 2025; Srivastava et al., 2026; Hunter and Martin, 2026), is fundamentally different from the problem studied here.

### Gradient basins are small

Gradient basins of peaks are much smaller than adaptive basins: Their scale is set by the size of the one-mutation neighborhood *n* = (*A*− 1)*L* of a genotype, rather than by the size *A*^*L*^ of the genotype space. Specifically, the average gradient basin size is 1 + *n* [Eq. (13)], and the maximal gradient basin size is 1 + *ne* [Eq. (15)]. These results are qualitatively consistent with Orr’s analysis of gradient walks, which shows that a typical walker finds a peak after *e*− 1 ≈ 1.72 steps (Orr, 2003). Moreover, our fitness-resolved analysis shows that, for large *n*, only genotypes with fitness very close to the maximum value *ω* = 1 have gradient basins that incorporate genotypes other than themselves.

The simple expression Eq. (15) for the mean size of the gradient basins is based on the fact that gradient basins are disjoint and exhaust the genotype space. Obviously, this remains true for other models of fitness landscapes. However, identifying the mean of the inverse number of peaks with the inverse of the mean requires additionally that the relative fluctuations in *N*_*p*_ vanish for large *L*, which does not necessarily hold for models of correlated fitness landscapes such as Kauffman’s NK-model (Durrett and Limic, 2003; Schmiegelt and Krug, 2014) or Fisher’s geometric model (Hwang et al., 2017; Park et al., 2020).

### Empirical patterns are consistent with our predictions

Large adaptive basins have been found in several recent empirical studies of fitness landscapes. Papkou et al. (2023) measured relative fitness of more than 260,000 variants of the *folA* gene in *Escherichia coli* in a mass-selection experiment under antibiotic stress, and determined the fitness peaks and the adaptive basins of the landscape. While the mean basin sizes for low and intermediate fitness peaks are below 0.02% and 5.7% of all genotypes, respectively, high fitness peaks were found to have a large mean adaptive basin size of around 69%. Our comparison focuses on the latter group of peaks, as in the HoC model the majority of peaks would be classified as high fitness peaks. Large adaptive basin sizes were also reported for fitness landscapes of bacterial transcrip-tion factors (Westmann et al., 2024; Antunes Westmann et al., 2024) and the *lac*-operator in *E. coli* (Chattopadhyay et al., 2025).

These studies corroborate two further key predictions of our analysis. First, the adaptive basin sizes of peaks are positively correlated with peak fitness. Specifically, Westmann et al. (2024) find a clean linear relationship between basin size and peak fitness rank, in agreement with the prediction of Eq. (5). Second, the empirical adaptive basins are dominated by indirect paths, with lengths that exceed the shortest distance between the corresponding genotypes by amounts between 10% and 80% (Papkou et al., 2023; Westmann et al., 2024; Antunes Westmann et al., 2024; Chattopadhyay et al., 2025). Of course, we do not mean to imply that the oversimplified random HoC model provides a quantitatively accurate representation of the empirical data sets. Nevertheless, the fact that the broad patterns of navigability predicted by the model are shared by the empirical landscapes suggests that they constitute robust structural features that apply across large classes of landscape models. In particular, the claim that the combination of high ruggedness and high navigability found in the empirical landscapes contradicts the predictions of classical models (Papkou et al., 2023) cannot be upheld in the light of our results.

### Limitations and open questions

Throughout this work we have adopted the simplest mathematical setup, assuming a homogeneous sequence space where all loci carry the same number of alleles and every allele is mutationally connected to every other allele. The framework of accessibility percolation on which our analysis is based can accommodate more general scenarios, where the number of alleles varies along the sequence and certain mutational transitions between alleles are forbidden. Under these modifications the threshold behavior for the probability of existence of accessible paths remains valid, but the threshold function takes a more complicated form and generally depends on the allelic compositions of the initial and the target genotype (Schmiegelt and Krug, 2023).

Another limitation to the applicability of our results arises from the fact that many empirical fitness landscapes contain missing genotypes, e.g., when nucleotide changes lead to a stop-codon or an otherwise nonfunctional sequence, or when the chosen fitness proxy signal is below the detection limit of the experiment (Franke et al., 2011; Papkou et al., 2023; Westmann et al., 2024). If individual genotypes are removed, the fitness graph loses the structure of a Cartesian power graph which is required for the approach of Schmiegelt and Krug (2023). Understanding the effect of missing genotypes on the navigability of fitness landscapes is an important problem for future research.

It would also be of interest to explore the statistics of adaptive basin sizes beyond the average size by computing higher moments. Accessible paths are strongly correlated. Determining the variance of the adaptive basin size would require the joint probability for two initial genotypes to have an accessible path to a target, which is not provided by Schmiegelt and Krug (2023) and would likely require a substantial extension of the underlying theory. The same problem arises in the analysis of gradient basins. However, as gradient basin sizes grow linearly rather than exponentially in *L*, using a tree-like approximation that neglects correlations between paths leads to a smaller relative error compared to the case of adaptive basins.

## Conclusion

Our theoretical and numerical results demonstrate that rough HoC fitness landscapes are highly navigable. Mean adaptive basins are of size *A*^*L*^(1−*β*^*\**^) and the fraction of accessible paths between any pair of genotypes approaches (1− *β*^*\**^)^2^ for large *L*, where *β*^*\**^ decreases with increasing number of alleles and tends to zero for large *A*. This phenomenon relies on the increasing connectivity of the genotype graph with increasing *A*, and is facilitated by indirect evolutionary paths which are significantly longer than the distance between the corresponding genotypes. These long paths are favored by simple sign epistasis and have been observed in empirical landscapes. However, this result needs to be interpreted with caution, as these long paths are rarely taken by adaptive walks.

Under greedy evolutionary dynamics, the picture changes fundamentally. Because gradient walks are strictly local and unable to utilize extended detours, the gradient basin of a peak is typically restricted to its one-mutant neighborhood and comprises a vanishingly small fraction of all genotypes for large *L*. Overall, low-dimensional metaphors, such as a physical mountainous terrain with peaks and valleys, fail to capture the importance of long paths for the navigability of highdimensional fitness landscapes under adaptive dynamics (Hernando et al., 2019), even if such metaphors retain some validity for gradient dynamics.

### Numerical methods

#### Adaptive Basins

We use depth-first search methods (Cormen et al., 2022) and simulate complete landscapes for the adaptive basin calculation, as computing a single adaptive basin requires checking a significant fraction of all genotypes for their membership. We sampled 10^7^ random genotypes for Fig.3**a** and 10^6^ peak genotypes for Fig.3**b**. Measurements within the same landscape are slightly correlated, as highly connected components tend to inflate these sizes globally. While this does not change the mean, it does affect the measurement error. Because the correlation is small, we allow for multiple (peak) genotypes to be sampled from the same landscape realization, but at most min 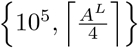 per realization.

For Fig.3**a** we sample at least 10^3^ and at most 10^7^ landscape realizations, and for Fig.3**b** at least 11 and at most 50005. Small landscape sizes, in terms of *L* and *A*, require the maximum number of landscape realizations, whereas large landscapes require the minimum.

### Gradient Basins

We implemented our own search algorithm for large *n* that does not make a tree-like approximation but respects the Hamming graph structure. This algorithm makes use of the local nature of gradient basins, as typical genotypes in a basin have Hamming distances *d* ≪ *L*, allowing for large *n* simulations. We need to check all genotypes at Hamming distance 2 from members of the gradient basin, leading to a complexity of roughly 𝒪 (*n*^3^).For each pair of *L* and *A*, we choose 100 evenly spaced fitness values between *ω*_2_ and *ω* = 1− 10^*−*6^, see Fig.4**a**. To ensure that the numerical mean and variance of the gradient basin size stabilizes, we simulate 2 ×10^4^ local landscape realizations for each fitness value.

## Code availability

Code used to produce the numerical results in the article will be made available upon publication.

## Acknowledgments

We thank Su-Chan Park for stimulating discussions and his eye for mathematical detail.

## Appendix A Accessibility Percolation

Let genotypes *τ*_*L*_ and *σ*_*L*_ have uniform (0, 1] fitness values 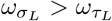 . Then Schmiegelt and Krug (2023) state that

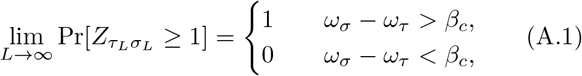

where 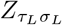 denotes the number of accessible paths from *τ*_*L*_ to *σ*_*L*_, and *ω*_*τ*_ and *ω*_*σ*_ are the limits of the sequences 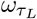and 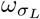. The threshold *β*_*c*_ is the unique root of the function Γ(*A, δ, β*) defined in Eq. (3). It is the exponential growth rate of the mean number of quasi accessible paths 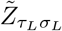. These are fitness-increasing paths that can, however, visit the same genotype multiple times by re-drawing its fitness value at each visit. Schmiegelt and Krug (2023) show that 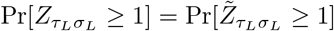.

The result (A.1) holds asymptotically for large *L*. At finite *L* the transition from low to high accessibility occurs over a fitness interval of order 𝒪 (1*/L*), whereas the threshold function *β*_*c*_ itself has a leading order correction of order 𝒪 (ln(*L*)*/L*) (Schmiegelt and Krug, 2023). Although the leading correction is explicitly computable, including only the𝒪 (ln(*L*)*/L*) term and no𝒪 (1*/L*) terms is not sufficient when comparing our results to numerical simulations. The largest system size we evaluate numerically is *L* = 28 for *A* = 2, where ln(*L*)*/L* ≈ 0.12 is not much larger than 1*/L* ≈ 0.036. For this reason we only use the asymptotic predictions of Schmiegelt and Krug (2023) for comparison with the numerical data in Fig. 3.

Schmiegelt and Krug (2023) reported analytical expressions for *β*_*c*_ and Γ for path endpoints at maximal distance *δ* = 1, but in this work expressions are needed 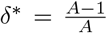 . Using *y* = *e*^*Aβ*^ as an ansatz, the problem reduces to finding the root of a polynomial in *y* of degree *A*. Analytical values of *β*_*c*_ for *A* = 2, 3 and 4 at *δ* = *δ*^*\**^ are shown in Table 1. For large *A* the ansatz 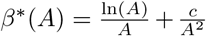 leads to Eq. (8) as a large *A* approximation, where terms including and higher than 𝒪 (*A*^*−*3^) are omitted. For large *A*, Eq. (8) differs by a factor of 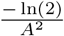 from the expression for *δ* = 1.

## Appendix B Adaptive basin size

This calculation is performed for fixed *A*. In the discrete distance picture, let *p*(*d, ω*) be the probability that there is an accessible path from a genotype at distance *d* from a target genotype with fitness *ω*. By linearity of expectation, the expected basin size of the target genotype is

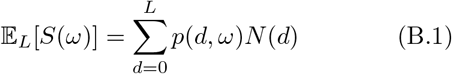

with *N* (*d*) from Eq. (4) and *p*(0, *ω*) = 1. This can be rewritten as

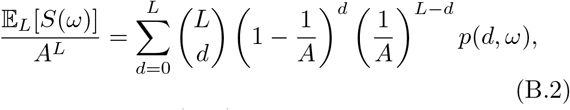

which shows that *p*(*d, ω*) is weighted by a binomial dis-tribution with parameter 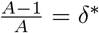. Increasing *L* results in most genotypes being at distance *Lδ*^*\**^. Switching to the re-scaled Hamming distance *δ* = *d/L* allows to approximate the binomial by a normal distribution *g*(*δ*) with mean *δ*^*\**^ and variance

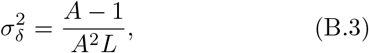

resulting in

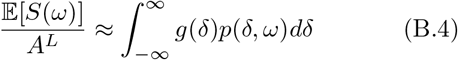

for large *L* as 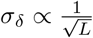 .

Next we calculate *p*(*δ, ω*). Let *τ*_*L*_ and *σ*_*L*_ be genotypes with fitness *ω*_*τ*_ and *ω*_*σ*_ = *ω* respectively, and re-scaled Hamming distance *δ*. Using Eq. (A.1), the probability that a fitness increasing path from *τ*_*L*_ to *σ*_*L*_ exists is

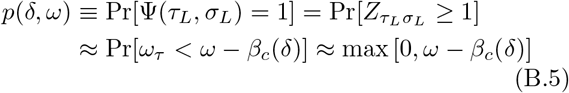

for large *L*. While *β*(*δ*) is defined by an implicit equation with no known analytical solution for general *A*, we can approximate *p*(*δ, ω*) around *δ*^*\**^. With *ϵ* = *δ* − *δ*^*\**^ we get 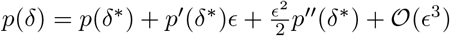. Usingthis approximation in Eq. (B.4) we obtain Eq. (5) with corrections of order 𝒪 (1*/L*), which are neglected for consistency with the approximations involved in arriving at (A.1).

## Appendix C Gradient Basins

In order to calculate the expected value of the size of the gradient basin of a target genotype with fitness *ω*, we first calculate the probability that *k* successive mutations yield the highest fitness gain in their respective neighborhood and form a strictly fitness increasing path. Consider a path of *k* strictly and maximally fitness increasing mutations, starting from a genotype *σ*_0_ with fitness *ω*_0_ to a genotype *σ*_*k*_ with fitness *ω*_*k*_, which belongs to the neighborhood of our target genotype *σ* with fitness *ω*. The fitness values *ω*_1_, *ω*_2_ … *ω*_*k*_ are the largest within their neighborhoods, and are therefore distributed according to *ω*_*i*_ ∼ max(*u*_1_, *u*_2_, …, *u*_*n−*1_) with *u*_*i*_ ∼ *U* (0, 1). This impliles that *ω*_*i*_ ∼ *f*_*ω*_(*x*) = (*n* − 1)*x*^*n−*2^. In the following we consider the *ω*_*i*_ to be independent, effectively approximating the fitness graph by a tree. Strictly speaking this is not true, as in a general Hamming graph genotypes share *A*−2 neighbors at distance one, but this effect is negligible for large *L*. Conditioning on the path to be fitness increasing, we get

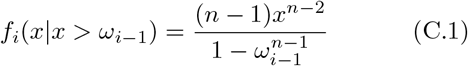

as the conditional density for the *ω*_*i*_ with ≥ *i* 1. Integrating over *k* steps and setting the boundary fitness values to *ω*_0_ and *ω* yields the expression

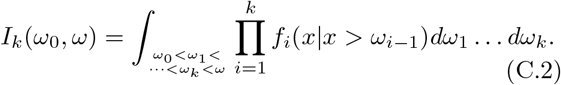

Substituting 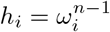results in

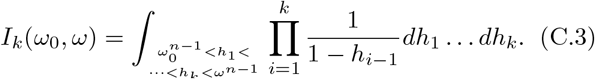

Since typically 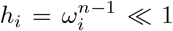 for large *n*, we can ap-proximate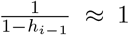. The integral then simply be-comes

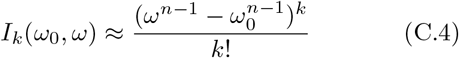

which is the volume of a *k*-dimensional simplex.

To complete the calculation, we need to account for the probability that the target genotype *σ* has the largest fitness among all direct neighbors of *σ*_*k*_. As we already conditioned on *ω*_*k*_ *< ω* in the simplex integral, this probability is given by *ω*^*n−*1^, and we arrive at the final result

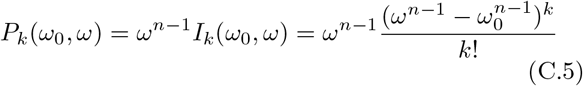

for the probability of the entire path.

From a starting genotype at distance *d* from the target, *d* of its neighbors are at distance *d*− 1, *d*(*A*− 1) at distance *d* and (*L*− *d*)(*A*− 1) at distance *d* + 1. Having fitter neighbors at distance *d* + 1 is therefore more likely than at distance *d* or *d*− 1, provided *d*≪ *L*. However, according to Eq. (C.4) every non-distance decreasing mutation incurs an additional factor ∝ *ω*^*n*^ for successive genotypes at distance *d* or ∝ *ω*^2*n*^ for genotypes at distance at *d* + 1. This implies that only direct (strictly distance-decreasing) mutational paths of genotypes contribute to the gradient basins. For a genotype at distance *l* from the target, a fraction *l/n* of mutations are distance decreasing. The restriction to direct paths therefore reduces the probability (C.5) by a factor

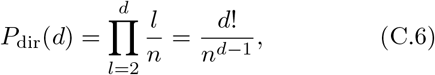

because the directness condition has already been enforced for the last step. Multiplying (C.5) with (C.6) and setting *k* = *d* − 1 results in the expression

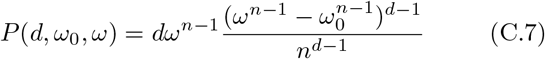

for the probability of a genotype of fitness *ω*_0_ at distance *d* to be part of the gradient basin of a target genotype with fitness *ω*.

As in the adaptive basin case we need sum over the whole genotype space, requiring an integration over *ω*_0_. However, for most genotypes 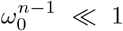 for large *n*, which implies that the dependence on the initial fitness can be omitted from Eq. (C.7). Moreover, large distances *d* are strongly suppressed in (C.7), which implies that we can approximate

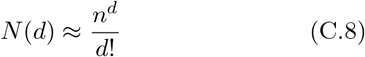

for *d* ≪ *L*. By linearity of expectation value we thus obtain

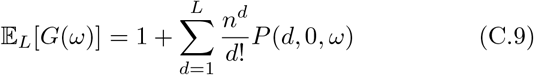

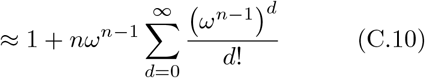

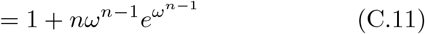

where we performed an index shift and took the limit *L*→ ∞, since contributions from *d > L* are negligible. Integrating Eq. (C.9) over the peak density and substituting *t* = *ω*^*n−*1^ recovers Eq. (13) asymptotically for large *n*,

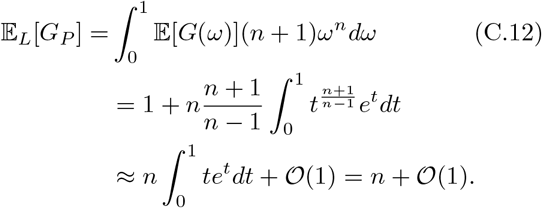

